# *fmr1* mutation interacts with sensory experience to alter the early development of behavior and sensory coding in zebrafish

**DOI:** 10.1101/2021.03.16.435742

**Authors:** Shuyu Zhu, Michael McCullough, Zac Pujic, Jordan Sibberas, Biao Sun, Bianca Bucknall, Lilach Avitan, Geoffrey J Goodhill

## Abstract

While Autism Spectrum Disorders (ASDs) are developmental in origin little is known about how they affect the early development of behavior and sensory coding, or how this is modulated by the sensory environment. The most common inherited form of autism is Fragile X syndrome, caused by a mutation in *FMR1*. Here we show that zebrafish *fmr1-/-* mutant larvae raised in a naturalistic visual environment display early deficits in hunting behavior, tectal map development, tectal network properties and decoding of spatial stimuli. However when given a choice they preferred an environment with reduced visual stimulation, and rearing them in this environment improved these metrics. Older *fmr1-/-* fish showed differences in social behavior, spending more time observing a conspecific, but responding more slowly to social cues. Together these results help reveal how *fmr1-/-* changes the early development of vertebrate brain function, and how manipulating the environment could potentially help reduce these changes.

## Introduction

Autism spectrum disorders (ASDs) are neurodevelopmental in origin. Increasing evidence suggests that a key way in which ASDs alter behavior and cognition is via altering the development of sensory processing [1]. While ASDs can be identified in humans as early as 6 months of age [2], little is known about how the early development of sensory neural processing is altered in ASDs.

Fragile X syndrome (FXS) is the most common single-gene cause of autism. It is due to a trinucleotide repeat expansion in the Fragile X mental retardation 1 (*FMR1*) gene, which leads to a lack of its product Fragile X mental retardation protein (FMRP). FMRP is highly expressed in neurons in the brain and regulates many aspects of brain development [3, 4, 5, 6]. Characteristics of the human FXS phenotype include low IQ, hyperactivity, attention deficits, and sensory deficits [1, 7, 8]. Changes in sensory processing are common in ASDs [1, 9, 10, 11, 12, 13, 14, 15, 16]. ASD individuals often display impaired adaptation to chronic sensory stimulation [17, 18, 19]. *Fmr1-/-* mice have circuit defects in the cortex [20, 21], larger networks of neurons that respond to sensory stimuli [22], and stronger motor responses and impaired adaptation to whisker stimulation [23]. However overall relatively little is known about how the early developmental trajectory of FXS affects behavior and sensory coding, and these are difficult questions to study in very young mammals.

In contrast the nervous system of zebrafish develops extremely rapidly, and by 5 dpf (days post-fertilization) larval zebrafish are already able to hunt fast-moving prey using only visual cues [24, 25, 26, 27]. This behavior relies on predictive models of target position [28]. Social behaviour begins to develop around 15 dpf and is again largely dependent on visual cues [29]. *nacre* zebrafish (which carry a mutation that affects pigment cells) are transparent at larval stages, and neural activity can be directly visualized non-invasively at large scale yet single-neuron resolution using transgenically encoded fluorescent calcium indicators in an intact and unanaethetised animal [30, 31]. Zebrafish have a strong genetic and physiological homology to mammals, and their affective, social and cognitive processes are analogous to those seen in rodents and humans [32]. However the effects of *fmr1-/-* mutation on the development of visually-driven behavior and associated neural coding remaing unknown.

While the environment has been hypothesized to play an important role in the expression of FXS, conflicting results have been obtained for how sensory experience affects the developmental trajectory of FXS mouse models. While [33] reported that environmental enrichment rescued some abnormalities, in contrast [34] found that enrichment was necessary for differences between the genotypes to be revealed. Since early zebrafish hunting and social behavior are highly visually driven and the complexity of visual stimulation can be easily manipulated, zebrafish provide a new opportunity to address the role of sensory experience in modulating the *fmr1-/-* phenotype.

Here we reveal that there is a delay in the early developmental trajectory of *fmr1-/-* compared to *fmr1+/-* zebrafish, reflected by less efficient and successful hunting behaviours at younger ages and delayed maturation of neural coding in the optic tectum. While these metrics normalised by 14 dpf, a longer-term effect of the mutation was revealed by altered social behavior at 28 dpf. However *fmr1-/-* fish preferred reduced sensory stimulation and, surprisingly, raising *fmr1-/-* fish in such an environment moved many of these of these metrics towards the *fmr1+/-* case. Together this work gives new insight into how *fmr1* mutation affects sensory development in the vertebrate brain, and provides evidence for an important impact of the environment on the development of FXS.

## Results

### *fmr1-/-* fish display craniofacial alterations

For this study we used the *fmr1-/-* knockout line generated from a TILLING screen by [35]. A characteristic feature of Fragile X syndrome is altered craniofacial structure, including an elongated face [36]. While craniofacial alterations were found in zebrafish *fmr1-/-* mutants generated using a morpholino knockdown approach [37], and subsequently in a CRISPR/Cas9 knockout [38], such changes were not originally reported in the knockout of [35]. We revisited this issue by crossing *fmr1+/-* with *fmr1-/-* fish to produce roughly equal numbers of *fmr1-/-* and *fmr1+/-* off-spring, performing Alcian blue staining at 3 developmental ages, and quantitatively comparing facial cartilage structure measurements (Fig. S1a,b). Canonical variate analysis [39] revealed differences in structure with both age (first canonical variable) and genotype (second canonical variable) (Fig. S1c). For the second canonical variable high weights were given for distances quantifying the length of the face (Fig. S1d), and at least two of these distances showed significant differences between genotypes at 9 and 14 dpf (Fig. S1e,f). In addition the angle of Meckel’s cartilage was significantly different between genotypes (Fig. S1g). These results confirm that craniofacial alterations analogous to human Fragile X syndrome occur in this *fmr1-/-* knockout, providing further support for this line as a relevant model system.

### Hunting is less successful in *fmr1-/-* fish

From 5 dpf zebrafish larvae start to hunt small, fast-moving prey such as *Paramecia*. This relies on precise sensorimotor coordination, and hunting success improves over development [27]. To test whether this behavior is altered by *fmr1* mutation, heterozygous and homozygous larvae were placed individually into small dishes with *Paramecia*, and hunting behavior was imaged for 10-15 min with a 500 fps camera. We imaged fish at 5, 8-9 and 13-14 dpf (henceforth referred to as 5, 9 and 14 dpf for brevity), and derived average values for hunting metrics across all events for each fish. Fish were genotyped after the experiment. To ensure we only included representative hunting behaviours, we used fish that had more than 7 hunting events across the entire duration of the hunting assay (10^th^ percentile of the distribution of number of events per fish; 9, 10, and 2 fish were rejected by this criterion for ages 5, 9, and 14 dpf respectively, leading to n = 21, 21, 10 for *fmr1-/-* and n = 20, 27, 11 for *fmr1+/-* for ages 5, 9 and 14 dpf respectively; different fish at each age).

*fmr1-/-* and *fmr1+/-* fish had similar gross motor function: fish length, speed, proportion of time stationary, number of bouts to strike, duration to strike and inter-bout interval were all indistinguishable between *fmr1-/-* fish and *fmr1+/-* fish (Fig. S2; these measures did though change with age, consistent with [27]). However *fmr1-/-* fish at 5 and 9 dpf were less successful at hunting than *fmr1+/-* fish, as evidenced by a lower hit rate (the fraction of successful prey captures out of all hunting events recorded per fish) (Fig. 1a), and higher abort rate (the fraction of abort events out of all hunting events recorded per fish, where an abort event means that the fish pursued the Paramecium of interest but aborted the pursuit and never struck at the prey) (Fig. 1b). 5- and 9-dpf *fmr1-/-* fish also showed a preference for hunting paramecia at more peripheral angles in the visual field (Fig. 1c) than *fmr1+/-* fish, as measured by the position of the target paramecium when eye convergence occurred, indicating the start of the hunting event (Fig. 1d). Together, these results demonstrate an initial delay in the development of effective hunting behavior *fmr1-/-* fish, and suggest an altered hunting strategy in these fish.

**Figure 1:**
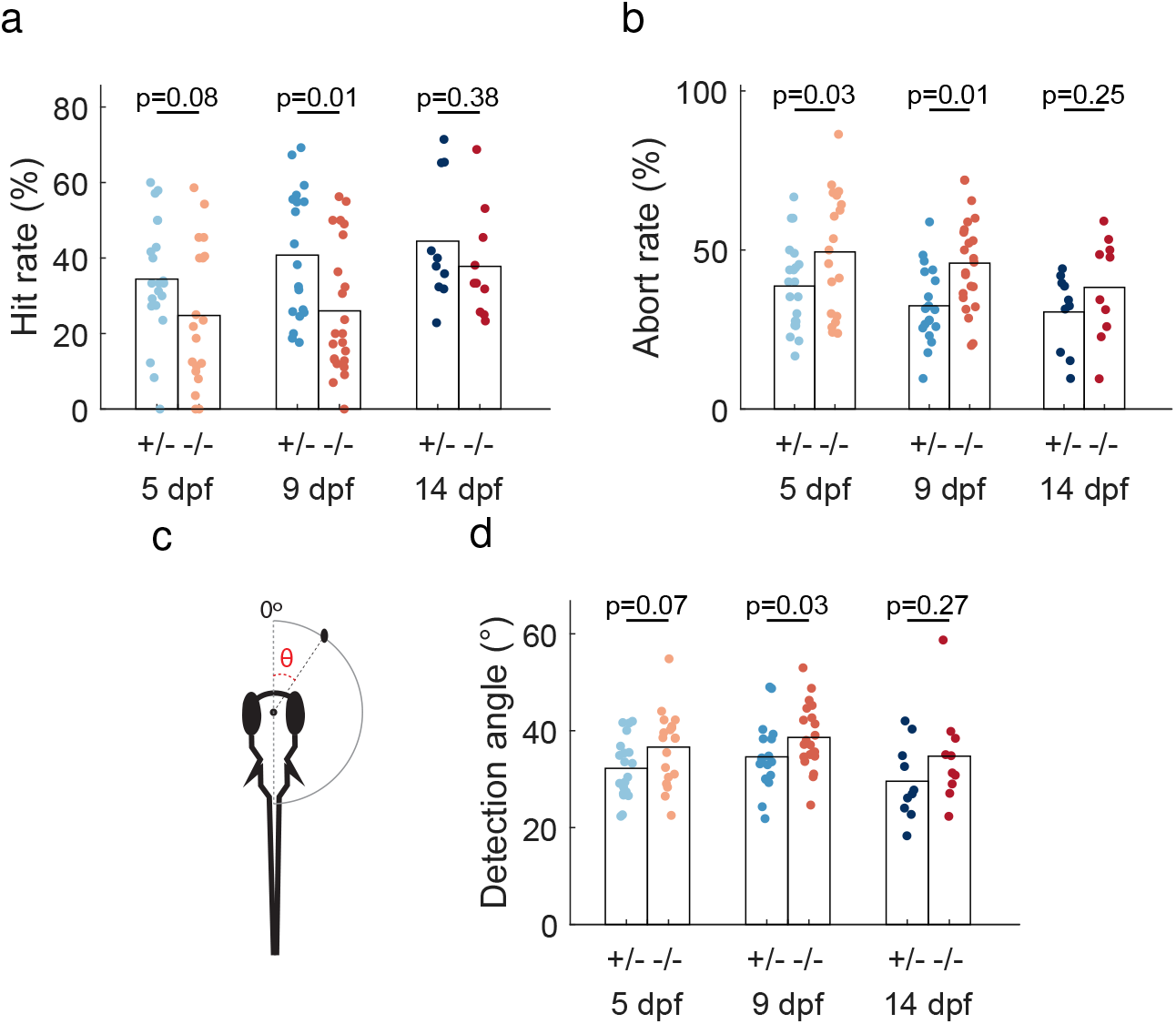
*fmr1-/-* fish show changes in hunting behavior. **a**. At 9 dpf *fmr1-/-* fish had a lower hit rate. **b**. At 5 and 9 dpf *fmr1-/-* fish had a higher abort rate. **c**. Prey angle was defined as the angle between the midline of the fish and the location of the paramecium prior to eye convergence (for detection angle) or after the first bout (after-bout angle). **d**. 9 dpf *fmr1-/-* fish responded to prey further towards the rear of the visual field.

### Stimulus-driven responses are slower to develop in *fmr1-/-* fish

In light of the changes in hunting in *fmr1-/-* fish observed above, we asked if *fmr1* mutation altered early development of spontaneous and evoked activity in the optic tectum, a brain region critical for successful hunting [40]. Fish aged at 5, 9 and 14 dpf (*fmr1-/-*, n = 10, 12, 6; *fmr1+/-*, n = 11, 12, 6 respectively) were embedded in low melting point agarose, and 2-photon imaging was used to record calcium signals from the tectum in a plane 70 µm below the skin [27]. Each fish was imaged first in the dark for 30 min of spontaneous activity (SA), followed by a 5 min adjustment period, and then in response to prey-like, 6°stationary spots at 9 positions in the visual field ranging from 45°to 165°in 15°increments. Each stimulus was presented for 1 s followed by a 19 s gap, with 20 repetitions of each stimulus in pseudo-random order. For some later analyses divided the data recorded for the stimulated period into activity from stimulus onset to 5 s post onset, (‘evoked activity’, EA) and activity from 15 s post-stimulus onset to the time of the next stimulus (‘spontaneous within evoked’, SE).

The tectum is topographically organised with the anterior portion responding to the frontal visual field, and the posterior portion responding to the rear visual field (Fig. 2a). However previous work with wild type fish has shown that the tectal representation of visual space at this tectal depth develops non-uniformly: responses are initially weaker and neural decoding worse in the anterior tectum, but by 13-15 dpf the representation has become uniform across the visual field [27]. We therefore asked if this developmental trajectory is altered in *fmr1-/-* fish. Responses in *fmr1-/-* fish were also topographically organised (Fig. 2b). However tectal development, as measured by the spatial uniformity of preferred stimuli, was initially delayed in *fmr1-/-* fish (Fig. 2c). The area under these curves was significantly smaller for *fmr1-/-* fish compared to *fmr1+/-* fish at 5 dpf, but equalised at later ages (Fig. 2d). The proportion of stimulus selective cells (those responsive to any stimulus) was lower for *fmr1-/-* compared to *fmr1+/-* fish at 9 dpf (Fig. 2e). Also, the proportion of tectal neurons responding to different visual angles was initially biased towards the rear visual field but became more evenly distributed over development for both *fmr1-/-* and *fmr1+/-* fish, similar to wild-type fish [27]. However at 5 dpf this bias was significantly more pronounced for *fmr1-/-* than *fmr1+/-* fish (Fig. 2f, 2g), again suggesting an intial developmental delay.

**Figure 2:**
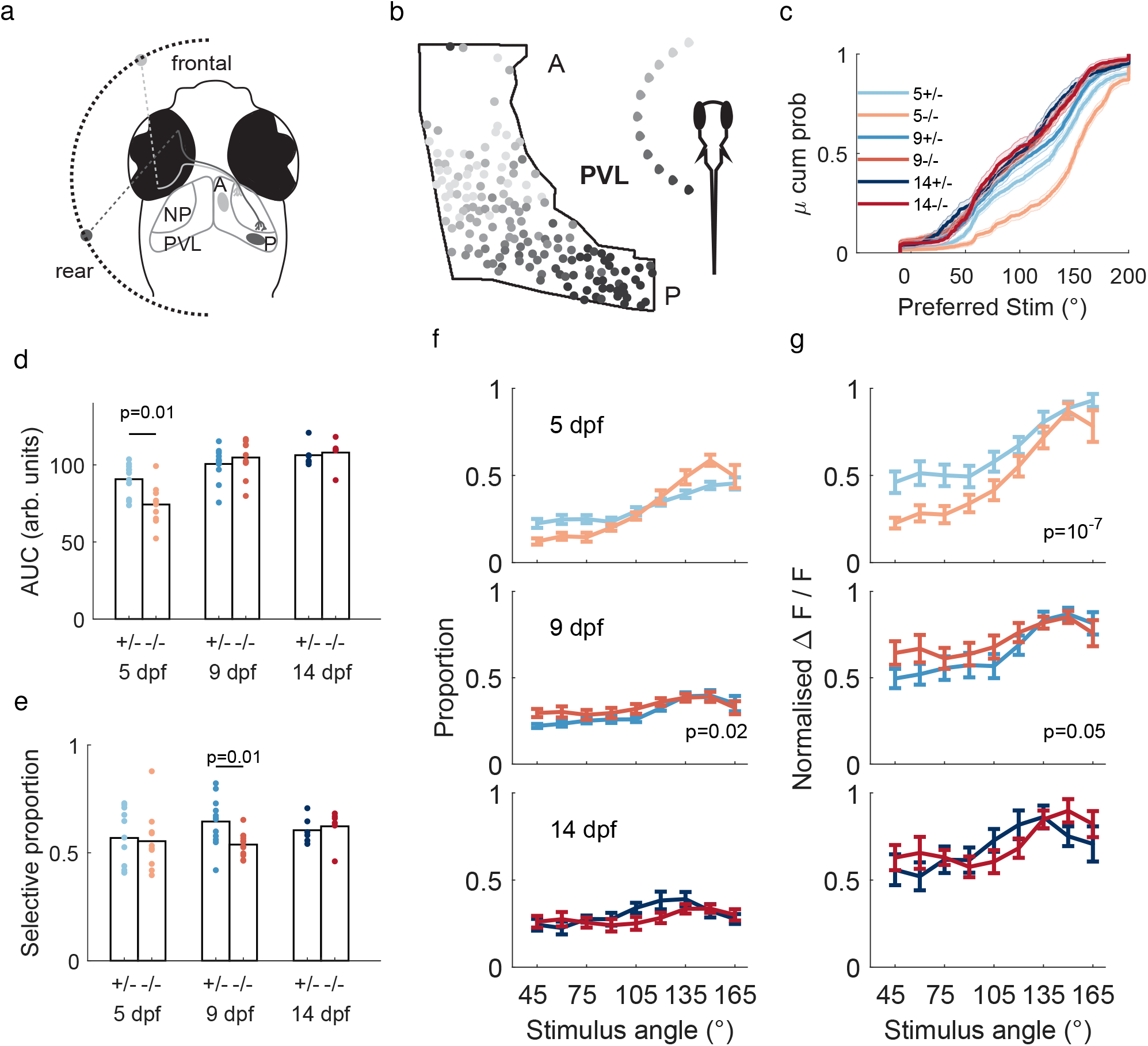
Tectal neurons in *fmr1-/-* fish show altered activity statistics. **a**. Schematic of the retinotectal projection in zebrafish. Retinal ganglion cells in the nasal part of the retina, representing the rear visual field, project to the posterior part of the tectum (dark grey). Retinal ganglion cells in the temporal part of the retina, representing the frontal visual field, project to the anterior part of the tectum (light grey). NP: neuropil; PVL: periventricular layer; A: anterior; P: posterior. **b**. Retinotectal projections are organised topographically in *fmr1-/-* fish (example 9-dpf fish). The stimulus position in the visual field to which each neuron in the PVL best responds is shown (see inset for grey-scale code). **c**. Cumulative distribution of preferred stimulus locations for both genotypes at 5, 9 and 14 dpf suggests a delay in 5-dpf *fmr1-/-* fish. **d**. Area under the curves in **c** shows that 5 dpf *fmr1-/-* fish had a less balanced representation of the visual field than 5-dpf *fmr1+/-* fish. **e**. Proportion of stimulus-selective neurons was lower in *fmr1-/-* fish at 9 dpf. **f**. Proportions of neurons responding to each stimulus angle were less balanced at 9 dpf for *fmr1-/-* fish. **g**. Responses to anterior stimuli were weaker in 5 dpf *fmr1-/-* fish. For **f**,**g** see panel c for color key. p-values indicate genotype effects using 2-way-ANOVA.

Thus at the level of individual neurons, *fmr1-/-* fish displayed an altered developmental trajectory of tectal spatial representation.

### Neural assemblies and neural coding are altered in *fmr1-/-* fish

Neural assemblies have been proposed to serve critical roles in neural computation [41]. We next identified tectal neural assemblies using the graph clustering algorithm introduced in [42] (Fig. 3a) and tested for alterations in assembly structure. For stimulus-evoked assemblies (EA) the number of neurons per assembly was greater for *fmr1-/-* than *fmr1+/-* fish at 9 dpf (Fig. 3b), suggesting higher excitability in *fmr1-/-* fish. However at 5 dpf assemblies in *fmr1-/-* fish were more compact, i.e. had a reduced span of their projection onto the AP axis of the tectum (Fig. 3c).

**Figure 3:**
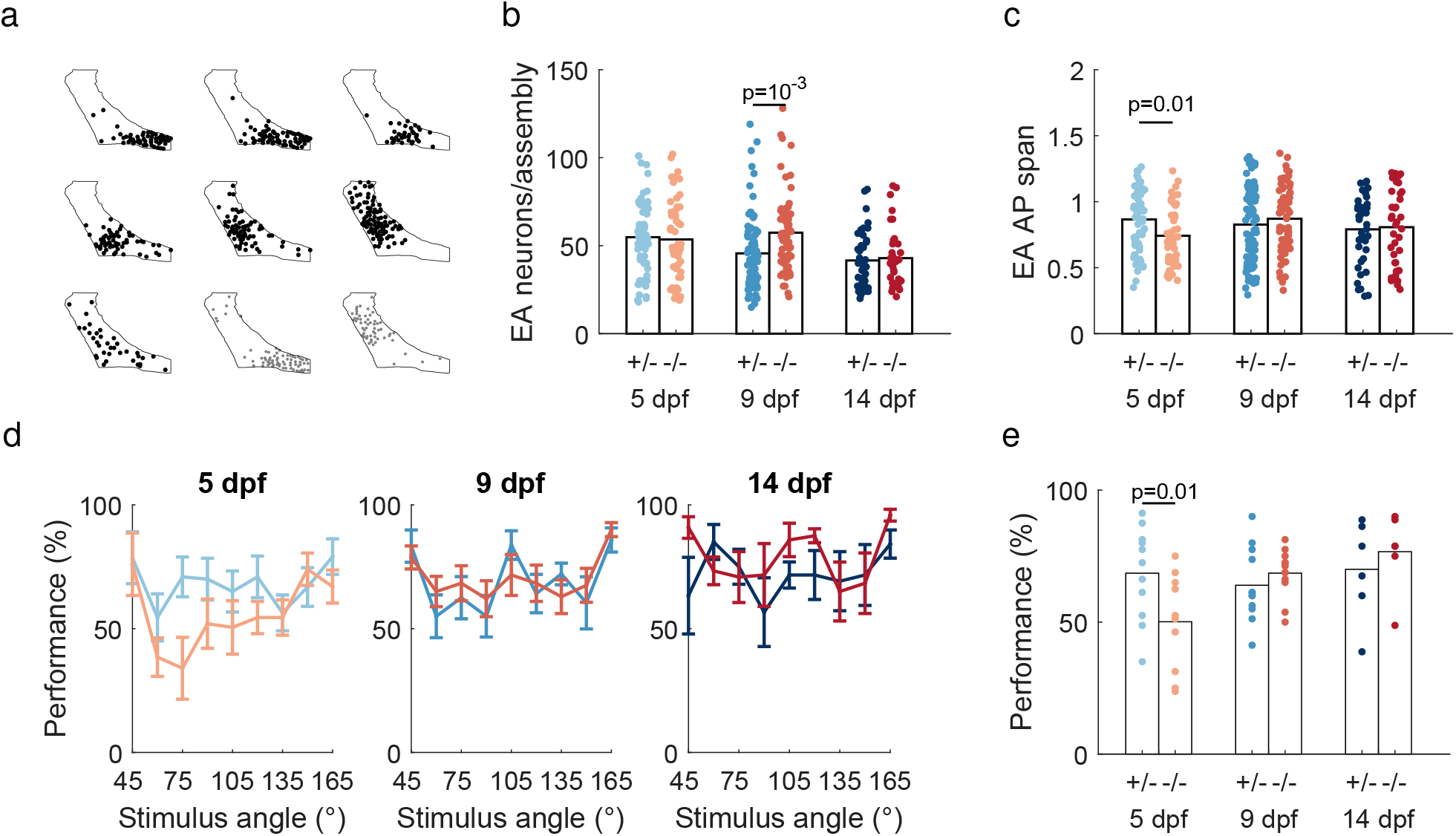
Neural assemblies and neural coding are altered in *fmr1-/-* fish. **a**. The assemblies detected in an example 8 dpf *fmr1-/-* fish drawn on the outline of the PVL. Black: EA assemblies. Gray: SA assemblies. **b**. At 9 dpf *fmr1-/-* fish had more neurons per EA assembly than *fmr1+/-* fish. **c**. At 5 dpf *fmr1-/-* fish had more compact assemblies. **d**. Comparison of decoder performance as a function of visual field position between genotypes at 5, 9 and 14 dpf. Color code as in earlier panels. **e**. Decoder performance averaged over frontal spots (up to 90°) was lower in *fmr1-/-* fish at 5 dpf.

These results suggest a delayed development of neural coding in the tectum. One measure of the quality of neural coding is decoding performance; in this case, how accurately stimulus position can be decoded from tectal activity. Decoding was worse for several visual field positions at 5 dpf for *fmr1-/-* fish, but this equalised over development (Fig. 3d-3e). Thus overall the developmental trajectory of tectal coding was altered in *fmr1-/-* fish, and displayed an initial delay relative to *fmr1+/-* fish.

### Correlation structures and synchronised activity patterns are altered in *fmr1-/-* fish

How are tectal network properties altered by *fmr1* mutation? During EA epochs short range correlations were higher for *fmr1-/-* fish (Fig. 4a), though similar for SA epochs (Fig. 4b). By thresholding the SA and EA correlation matrices and determining their degree of similarity (Hamming distance), we found that these matrices were less similar for 5 dpf *fmr1-/-* fish (Fig. 4c). At 9 dpf there was an increase in coactivity level (mean number of neurons active together) in *fmr1-/-* fish for EA epochs (Fig. 4d). At 9 dpf EA epochs for *fmr1-/-* fish had higher dimensionality, as measured by the Participation Ratio [43] (Fig. 4e). The residuals for both SA and SE patterns when projected onto the EA space (see Additional methods) were larger in *fmr1-/-* fish at 9 dpf (Fig. 4f), suggesting EA patterns in these fish were geometrically less similar to SA patterns than in *fmr1+/-* fish. However, we did not observe such differences at 5 or 14 dpf (Fig. S3).

**Figure 4:**
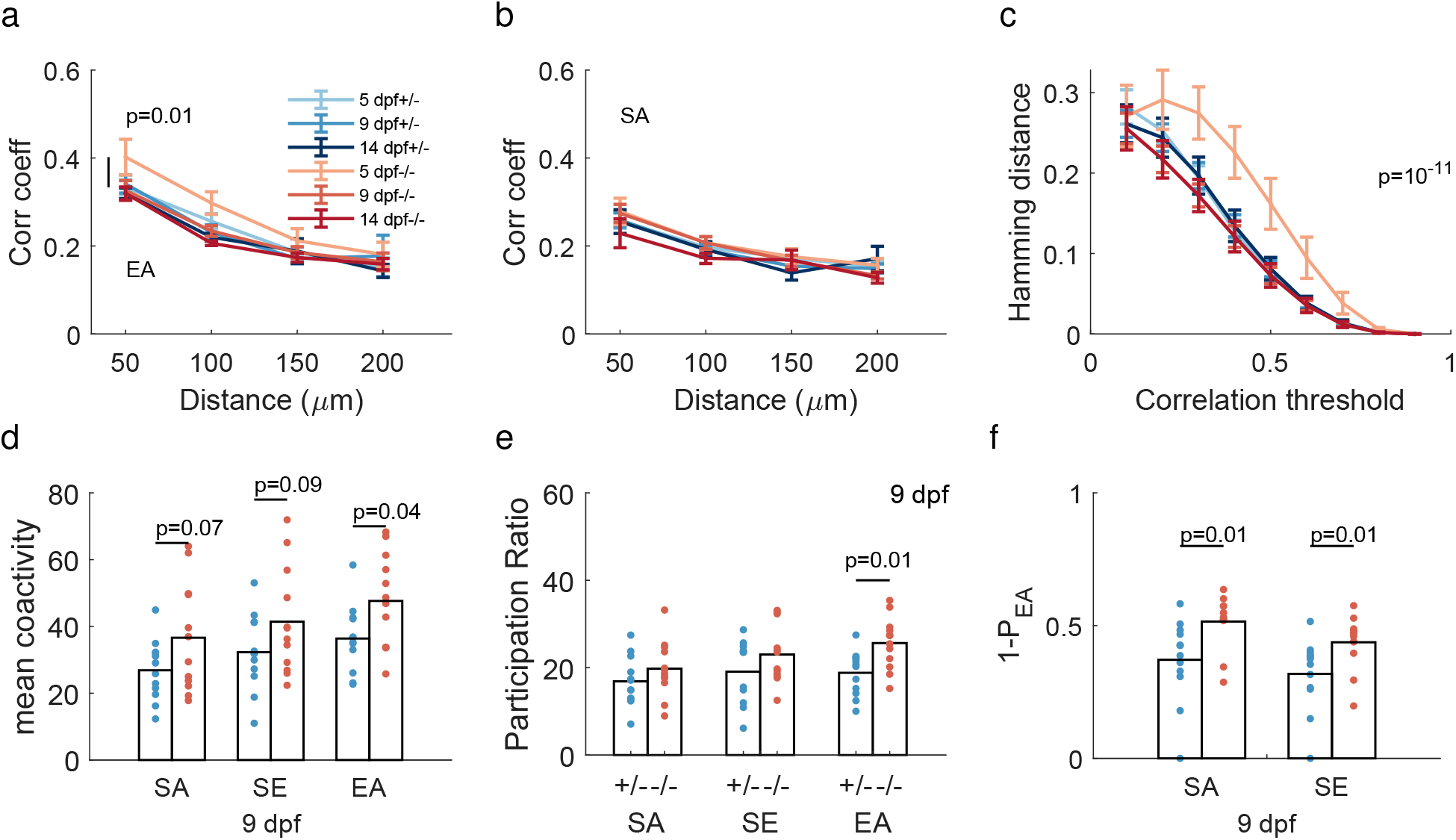
Network properties are altered in *fmr1-/-* fish. **a**. At 5 dpf EA correlations were greater at short range (*<*50 µm) for *fmr1-/-* fish. **b**. SA correlations were similar between genotypes. **c**. The similarity between EA and SA correlation structures was lower at 5 dpf for *fmr1-/-* fish (color scheme as in a). **d**. The number of coactive neurons during EA at 9 dpf was higher for *fmr1-/-* fish. **e**. The dimensionality of evoked activity at 9 dpf was higher for *fmr1-/-* fish, as measured by the participation ratio. **f**. The residuals of the projections of SA and SE onto the EA space were larger in 9-dpf *fmr1-/-* fish.

Thus at early ages compared to *fmr1+/-* fish, *fmr1-/-* fish had higher correlations between neurons, decreased similarity between evoked and spontaneous activity patterns, higher coactivity levels and higher-dimensional activity, consistent with increased excitability. However these properties had mostly equalised by 14 dpf, suggesting a transient period of disorder in network properties during development.

### Reduced sensory stimulation during development improves outcomes for *fmr1-/-* fish

For the experiments described thus far the fish were raised in petri dishes placed on a gravel substrate [44] (see Additional methods), which is a more natural visual environment than featureless petri dishes, and is indeed preferred by adult wild-type fish [45]. However humans with ASDs often experience sensory over-responsitivity to normal sensory environments, sometimes accompanied by aversive behaviours [46]. We therefore wondered whether *fmr1-/-* larvae would prefer an environment with reduced sensory stimulation, and whether rearing in such an environment would change developmental outcomes for these fish.

First we compared free-swimming behavior (no prey items) for *fmr1-/-* and WT fish at 8-9 dpf in 85 mm dishes, where half of each dish had an image of a gravel substrate on the bottom and the other half was featureless (uniform brightness equal to the mean brightness of the gravel half of the dish) (Fig. S4a). WT fish displayed no preference for either side of the dish. However *fmr1-/-* fish spent significantly more time on the featureless side of the dish (Fig. 5a), consistent with the hypothesis of an active avoidance of sensory stimulation. This was true both for fish raised to that point on gravel, and fish raised in a featureless environment (Fig. S4b).

**Figure 5:**
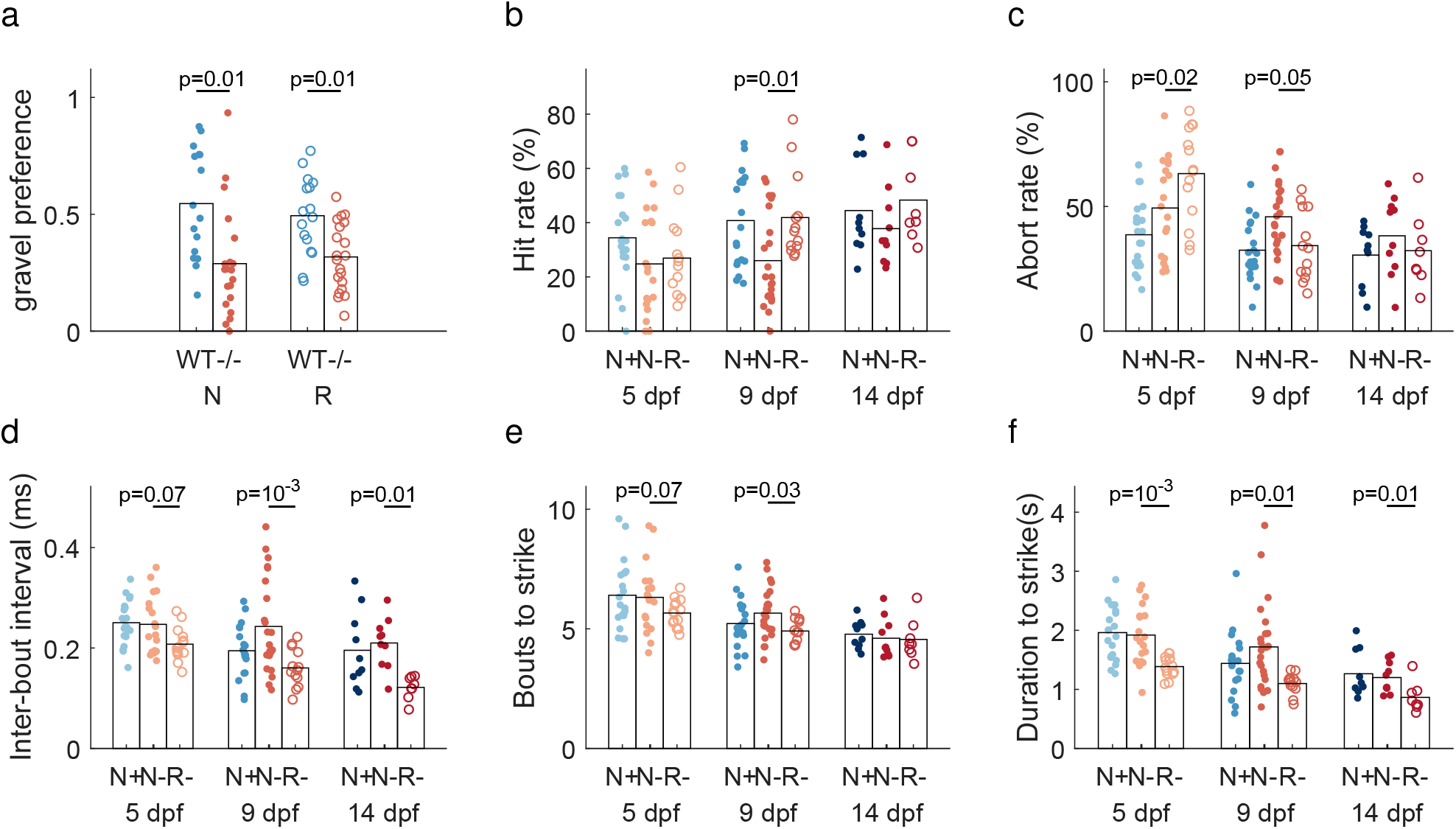
Reduced sensory stimulation improves hunting behaviours in *fmr1-/-* fish. **a**. *fmr1-/-* fish preferred a featureless to gravel environment, but *fmr1+/-* fish had no preference. N: fish reared under naturalistic conditions; R: fish reared under reduced stimulation (featureless) conditions. p-values shown are 2-sample t-test. Results for 1-sample t-tests comparing each sample with 0.5 were 0.4 (N) and 0.9 (R) for WT, and 0.0007 (N) and 0.00002 (R) for *fmr1-/-*. **b-f**. Terminology: R-, *fmr1-/-* fish raised with reduced sensory stimulation; N-, *fmr1-/-* fish raised under naturalistic conditions; N+, *fmr1+/-* fish reared under naturalistic conditions (shown for comparison, same data as Figs 1-4). **b**. Hit ratio was higher for *fmr1-/-*(R) than *fmr1-/-*(N) fish at 9 dpf, towards the *fmr1+/-*(N) case. **c**. Abort rate was greater for *fmr1-/-*(R) than *fmr1-/-*(N) fish at 5 dpf, but less at 9 dpf, towards the *fmr1+/-*(N) case. **d-f**. *fmr1-/-*(R) fish were more efficient in hunting than *fmr1-/-*(N) fish with shorter inter-bout interval (9 and 14 dpf), less bouts to stike (9 dpf) and shorter duration to strike (all ages).

Next, we compared our original cohort of *fmr1-/-* fish raised on gravel (now termed *fmr1- /-*(N), for ‘naturalistic stimulation’) with a new cohort of *fmr1-/-* fish raised in featureless dishes (termed *fmr1-/-*(R), for ‘reduced stimulation’), in order to determine whether the sensory environment could affect the expression of the *fmr1-/-* phenotype (n = 9, 14, 6 for 5, 9, 14 dpf respectively). Statistical comparisons are presented between *fmr1-/-*(N) and *fmr1-/-*(R) fish, but the data discussed earlier for *fmr1+/-*(N) fish is also shown again for comparison.

When assessed using the same featureless chambers as before, hunting success (hit ratio) was significantly improved at 9 dpf for *fmr1-/-*(R) compared to *fmr1-/-*(N) fish (Fig. 5b). This was primarily driven by a decrease in the abort ratio for *fmr1-/-*(R) fish (Fig. 5c). However at 5 dpf the abort rate for *fmr1-/-*(R) fish was higher than *fmr1-/-*(N) fish, despite there being no difference in hit rate, suggesting that *fmr1-/-*(R) fish had difficulty sustaining hunting events at this early age. We found that across a range of ages *fmr1-/-*(R) fish were more efficient at hunting, as measured by inter-bout interval during a hunting sequence (Fig. 5d), number of bouts prior to a strike (Fig. 5e), and duration to strike (Fig. 5f).

Reduced sensory stimulation also altered tectal responses in *fmr1-/-* fish. At 9 dpf neurons in *fmr1-/-*(R) fish were less excitable (Fig. 6a) with smaller tuning widths (Fig. 6b). 9 dpf *fmr1-/-*(R) fish also had fewer neurons per EA and SA assembly than *fmr1-/-*(N) fish (Fig. 6c,6d). *fmr1-/-*(R) fish had less compact EA assemblies at 5-dpf *fmr1-/-*(R) fish compared to *fmr1-/-*(N) fish (Fig. 6e). At 9 dpf coactivity levels in *fmr1-/-*(R) fish were lower than *fmr1-/-*(N) fish during EA epochs (Fig. 6f). We also found that both SA and SE patterns in *fmr1-/-*(R) fish were geometrically more similar at 9 dpf to EA patterns compared to *fmr1-/-*(N) fish (Fig. 6g). For all these metrics the *fmr1-/-*(R) fish were closer to the *fmr1+/-*(N) fish than were *fmr1-/-*(N) fish. Thus reduced sensory stimulation during development reduced the impact of the *fmr1* mutation.

**Figure 6:**
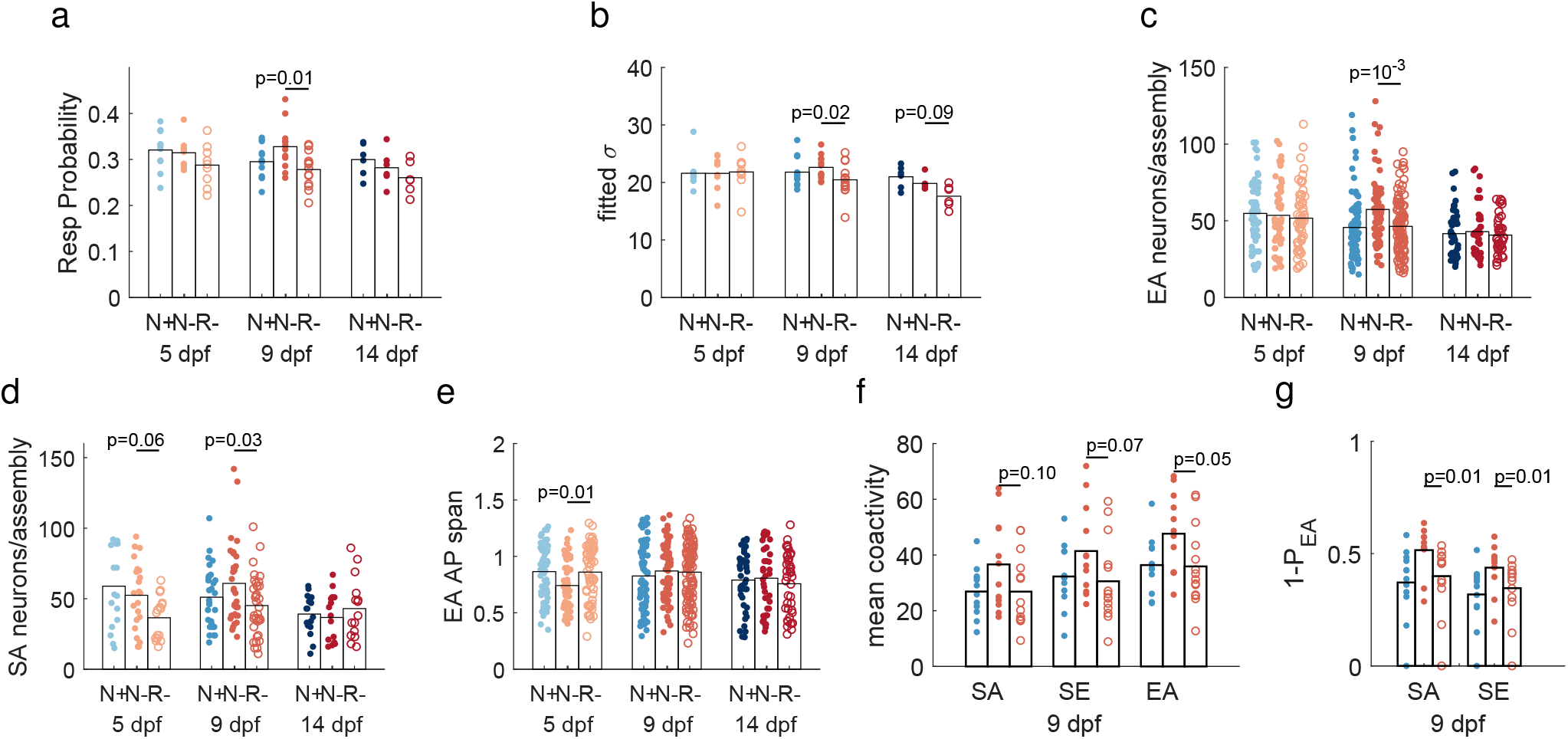
Reduced sensory stimulation in *fmr1-/-* fish moves tectal activity closer to the *fmr1+/-*(N) case. **a**. At 9 dpf neuron response probability was lower for *fmr1-/-*(R) fish, towards the *fmr1+/-*(N) case. **b**. At 9 dpf neurons in *fmr1-/-*(R) fish had smaller tuning width compared to *fmr1-/-*(N) fish, towards the *fmr1+/-*(N) case. **c-d**. At 9 dpf *fmr1-/-*(R) fish had less neurons per assembly for both EA (c) and SA (d) assemblies, towards the *fmr1+/-*(N) case. **e**. EA assembly members spanned more of the AP axis in *fmr1-/-*(R) fish at 5 dpf, towards the *fmr1+/-*(N) case. **f**. 9 dpf *fmr1-/-*(R) fish had lower coactivity levels than *fmr1-/-*(N) fish for EA epochs, towards the *fmr1+/-*(N) case. **g**. When projected onto the subspace of EA patterns, SA patterns of the 9 dpf *fmr1-/-*(R) fish had smaller residuals, towards the *fmr1+/-*(N) case.

### Social behaviour is altered in *fmr1-/-* fish

For many of the metrics examined above, by 14 dpf *fmr1-/-* fish are indistinguishable from *fmr1+/-* fish. Does this mean that the effects of *fmr1* mutation in zebrafish are only transient? A key behavior that emerges at later ages is social interaction. We therefore asked whether there are any differences in social behavior between *fmr1-/-* and WT fish, at both 13-14 dpf and 26-28 dpf (WT: n = 36, 88, *fmr1-/-*: n = 48, 80 respectively; for simplicity we will refer to these as just 14 and 28 dpf respectively; these fish were raised in 1 L tanks in the University of Queensland’s central aquarium). For these experiments we used a U-shaped behavioral chamber similar to that of [47] (Fig. 7a,b), and compared how the movements of *fmr1-/-* versus WT test fish were affected by the presence of a WT cue fish in one arm of the chamber over an imaging time of 30 min. To avoid potential effects on social behaviour caused by differences in physical appearance, both WT and *fmr1-/-* fish were in nacre background and the cue fish was size matched to the test fish.

**Figure 7:**
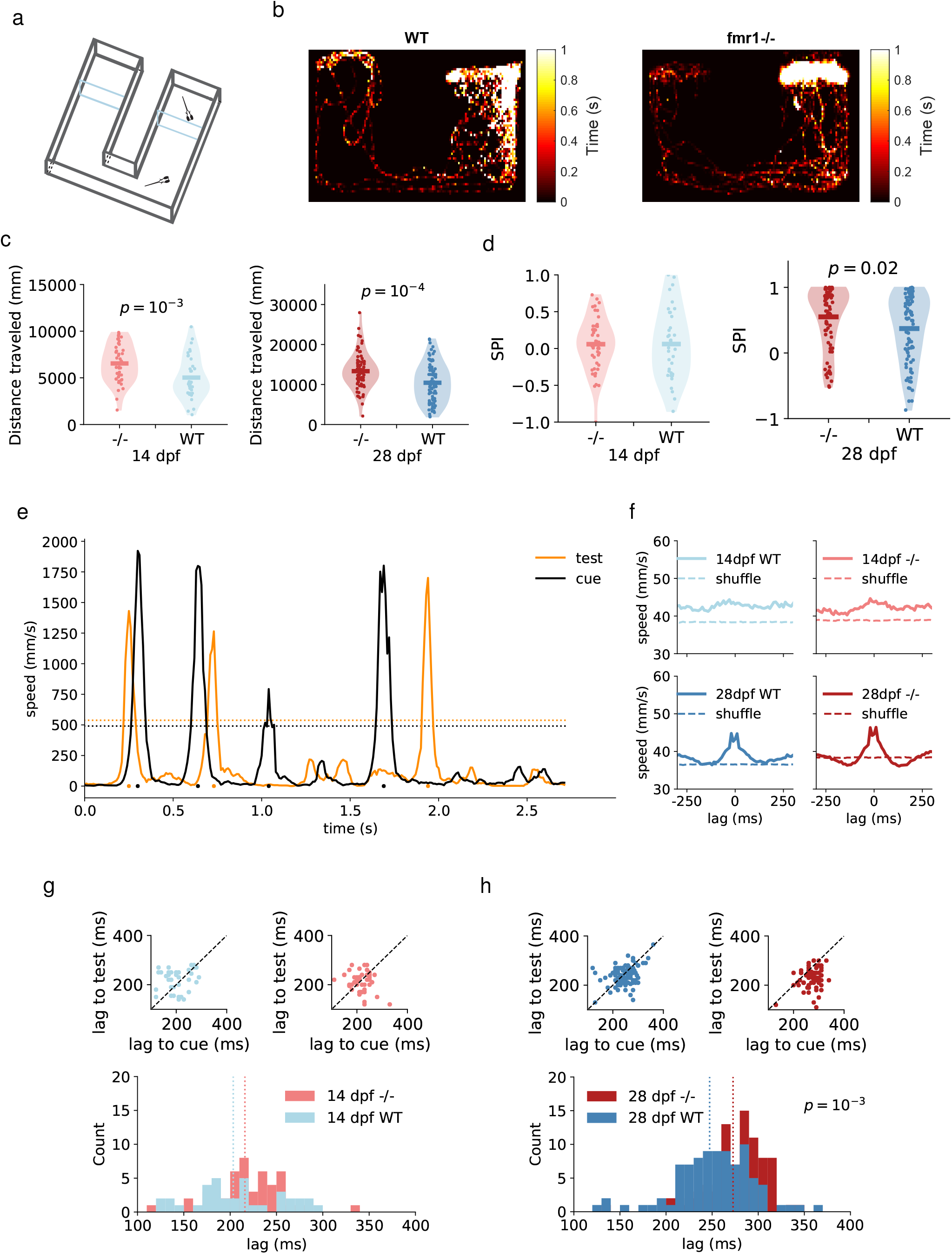
*fmr1-/-* fish display altered social behavior. **a**. Illustration of the chamber used for the social assay. **b**. Example heat maps of the position of the test fish over 30 min (28 dpf, WT SPI: 0.75; *fmr1-/-* SPI: 0.84). **c**. Total distance traveled was greater for *fmr1-/-* than WT fish at both 14 and 28 dpf. **d**. At 28 dpf social preference index (SPI) was higher for *fmr1-/-* fish. **e**. An example temporal segment of fish speed illustrating that the fish respond to each other’s movements, and that either fish can lead. Dashed line represents significant motion threshold level. Each dot indicates a significant movement peak time. **f**. Averaged motion signal for 200 ms each side of movement peaks confirmed coordinated movements at 28 but not 14 dpf. **g - h**. Average movement lag was longer for *fmr1-/-* fish at 28 dpf.

At both 14 and 28 dpf, *fmr1-/-* fish travelled a greater distance in the chamber than WT fish (Fig. 7c). This is consistent with hyperactivity of *fmr1-/-* fish as reported previously [48, 49]. As an initial measure of social interaction we calculated the social preference index (SPI) as in [47], which measures the proportion of time the fish spends in the arm of the chamber containing the cue fish versus the empty arm. Neither genotype displayed a preference between arms at 14 dpf (Fig. 7d), but by 28 dpf both genotypes showed a preference for the arm containing the cue fish. Surprisingly however, at 28 dpf *fmr1-/-* fish had a stronger preference than WT fish for the arm containing the cue fish, suggesting a greater desire for social interaction (Fig. 7d).

When cue and test 28-dpf fish could see each other they tended to respond to each other’s movements, with sometimes the test fish leading and sometimes the cue fish leading (Fig. 7e). This behavior was not present at 14 dpf (Fig. 7f, 7g). However by 28 dpf *fmr1-/-* fish, unlike WT fish, showed a clear asymmetry between their behavior and that of the cue fish. In particular, *fmr1-/-* fish took on average 26 ms longer than WT fish to respond to movements of the test fish (Fig. 7h). Thus it appears that, although *fmr1-/-* fish have greater desire for social interaction than WT fish, they interact less effectively.

## Discussion

Previous studies of zebrafish mutant for *fmr1* have shown a variety of phenotypic effects. Using a morpholino approach [37] reported changes in craniofacial structure and increased axonal branching during development. The initial description of the knockout line used in the present work did not find similar changes [35], which has led to doubts about the relevance of this line for studying FXS [38]. However [35] did not report quantitative results for craniofacial structure. Our more detailed and rigorous analysis demonstrates that craniofacial abnormalities do indeed exist in this line (Fig. S1). Using this line [50] showed changes in open-field behavior in adult *fmr1-/-* fish, [51] showed increased axonal branching early in development, and [52] showed abnormal auditory processing. Using adults from a different *fmr1* knockout line, [48] showed changes in exploratory behavior, avoidance learning, long-term potentiation and long-term depression. Using a *fmr1* knockout generated via CRISPR/Cas9, [38] showed that 5 dpf fish had craniofacial changes, hyperactivity, and changes in response to light stimulation. Here we have significantly extended these previous analyses of the *fmr1* knockout by examining hunting and social behavior, tectal coding, how these change across development, and how the visual environment can alter the expression of the *fmr1* knockout phenotype.

In terms of tectal activity we found an altered developmental trajectory of tectal spatial representation and tectal coding in *fmr1-/-* fish, including higher correlations and coactivity levels at younger ages. Many of these changes mirror those seen previously in *Fmr1-/-* mouse cortex [20, 53], supporting the relevance of zebrafish model. These include larger short-range neuron-neuron correlations at young ages, and larger numbers of neurons recruited to peaks of synchrony (analogous to our neural assemblies). A leading hypothesis for the underlying cause of some of these changes is an increase in neural excitatation (E) relative to inhibition (I), i.e. E-I balance [54]. Supporting this, inhibitory interneurons have been implicated in network dysfunction in FXS [55, 56, 57, 58]. A recent suggestion is that E-I balance changes are in fact compensatory in ASDs, helping to restore the system to a normal operating point [59]. Inhibitory neurons in zebrafish tectum have been identified using a variety of molecular techniques. For instance, [60] found that almost all *dlx5*-positive neurons in the tectum are GABAergic, and that this population comprises 5 - 10% of all tectal neurons. While alterations in E-I balance in *fmr1- /-* zebrafish remain to be investigated, an intriguing hypothesis raised by our work is that any such changes are modulated by the environment in which the animals are raised.

Our behavioral data shows that, at younger ages, *fmr1-/-* fish are worse hunters than *fmr1+/-* fish under naturalistic rearing conditions. Given the changes we observed in tectal activity, this is consistent with findings from mice [61, 62] and humans [2, 9, 11, 12, 14, 63] that *fmr1* mutation introduces low-level visual deficits. However, according to some metrics, *fmr1-/-* fish raised with reduced sensory stimulation were better at prey capture than *fmr1-/-* raised under naturalistic conditions. It should be noted though that all prey-capture assays were perfomed in relatively featureless dishes, a similar visual environment to the reduced sensory stimulation rearing case. This could potentially place fish raised under naturalistic conditions at a disadvantage in our prey capture assay, since they have adapted to hunting in a richer visual environment than the reduced stimulation case. This would be potentially analogous to recent reports that whether zebrafish first experience dry or live food influences their subsequent behavior and brain development [64, 65].

For efficiency our primary comparisons were between *fmr1-/-* and *fmr1+/-* fish, both generated from crossing *fmr1-/-* and *fmr1+/-* fish. For our neural imaging experiments we could only examine one fish per day, and the genotype could only be determined after each imaging or behavioral experiment using PCR. Thus crossing *fmr1-/-* with *fmr1+/+* fish to additionally compare *fmr1-/-* and *fmr1+/-* with *fmr1+/+* would have required twice as many experiments to obtain the same n values per group. Whether a comparison of *fmr1-/-* and *fmr1+/+* fish would yield stronger or additional phenotypic differences according to the measures we have examined remains a question for future work; however this caveat does not weaken our conclusions regarding differences we have observed between *fmr1-/-* and *fmr1+/-* fish.

A common symptom of human FXS is sensory hypersensitivity, which can lead to sensory defensiveness [66]. Consistent with visual hypersensitivity we found that *fmr1-/-* fish, unlike *fmr1+/-* fish, preferred to swim in an environment with reduced visual stimulation compared to naturalistic conditions. This is analogous to findings of tactile defensiveness in *Fmr1-/-* mice [23]. Furthermore, tectal neurons in our *fmr1-/-* fish showed trends towards higher response probability, and a larger number of neurons per assembly for evoked activity. However, reducing visual stimulation during development moved several metrics of behavior and tectal coding closer to those of *fmr1+/-* raised in a naturalistic environment. A comparison can be made with studies of *Fmr1-/-* mice examining the effects of environmental enrichment (EE) (e.g. running wheels and toys). [33] showed that EE largely rescued symptoms of hyperactivity, open-field exploration, habituation and changes in dendritic structure compared to mice reared in the normal lab environment [33], and a subsequent study showed restoration of long-term potentiation in prefrontal cortex to wild type levels [67]. While this would appear to conflict with our results for zebrafish, more recent work found that hippocampal spine morphology was more different between *Fmr1-/-* and WT mice after EE [34]. These authors suggested that EE allows for the impact of loss of *Fmr1* to be more fully expressed, which is more consistent with our findings. Overall our work suggests an important role for the sensory environment in modulating the effects of loss of *fmr1-/-*, with potential implications for therapies.

Many of the differences in prey capture and neural properties we observed in *fmr1-/-* fish occurred at 9 dpf. A previous study of the development of spontaneous neural activity in zebrafish tectum suggested that major reorganisations of tectal networks may be occurring just before this, at 5-6 dpf [42]. Assuming that lack of *fmr1* takes some time to manifest, this would be consistent with observing changes slightly later. Interestingly many of these properties became indistinguishable between genotypes at 14 dpf. However this does not mean that the system had necessarily returned to a normal developmental trajectory by this age. First, we observed changes in social behavior at 28 dpf, even though these were not apparent at 14 dpf. Second, it has been argued that a misregulation of critical periods can have very long-lasting effects [68]. The loss of a particular gene product can result in compensatory regulation of other genes, but this compensation takes time, meaning that critical windows for time-sensitive developmental events may be missed. This hypothesis explains why overall the system may ultimately not function normally, even though some aspects which are initially delayed eventually catch up.

We found that by 28 dpf *fmr1-/-* fish display a greater preference for social interaction with a cue fish than *fmr+/-* fish. This is initially surprising, given the well-documented tendency in ASDs in general for reduced social interaction [69]. However, recent work suggests that FXS may diverge from typical ASDs in this regard. In particular, [70] found in an eye-gaze paradigm that individuals with FXS did not show the large reductions in social interest characteristic of idiopathic ASDs. On the other hand, we also found a reduced effectiveness of social interaction in *fmr1-/-* fish, in terms of a slower response to movements of the cue fish. This could potentally be simply a motor deficit, but we found no direct evidence for motor deficits in *fmr1-/-* fish in the prey-capture assay. The altered interaction efficiency observed here is consistent with a recent report of deficits in imitating conspecific behaviour in *Fmr1-/-* mice [71]. A more likely explanation is an alteration in information processing in the networks underlying social interaction [72, 73], and analysing these in *fmr1-/-* fish is an interesting direction for future work.

Together our results reveal many previously unknown differences in natural behavior in *fmr1-/-* fish, and neural bases for these behavioral changes in terms of altered neural coding. The changes in the developmental trajectory of *fmr1-/-* fish depending on the complexity of the sensory environment, with a less complex environment leading to better outcomes, offers a new direction for future work, potentially leading to novel concepts for therapeutic intervention. Overall, our work suggests new avenues for revealing the developmental alterations of neural systems in neurodevelopmental disorders.

## Materials and Methods

### Zebrafish

All procedures were performed with the approval of The University of Queensland Animal Ethics Committee. Fish with the *fmr1*^hu2787^ mutation were originally generated by the Ketting laboratory [35], and obtained for this study from the Sirotkin laboratory (State University of New York). We first in-crossed the mutant line to generate nacre *fmr1*^hu2787^ mutants. For calcium imaging and hunting assay experiments these nacre *fmr1*^hu2787^ mutants were crossed with nacre zebrafish expressing the transgene *HuC:H2B-GCaMP6s* to give pan-neuronal expression of nuclear-localised GCaMP6s calcium indicator. *fmr1+/-* were then crossed with *fmr1-/-* fish (with no consistent relationship between the genotype and the sex of the parent) to produce roughly equal numbers of *fmr1+/-* and *fmr1-/-* offspring. For social behaviour assays nacre *fmr1*^hu2787^ mutants were used.

Fish embryos were raised in E3 medium (5mM NaCl, 0.17mM KCl, 0.33mM CaCl_2_, 0.33mM MgCl_2_) at 28.5° C on a 14/10 h light/dark cycle. For the data in Figs 1-6 Fish were kept in small groups in 100 mm petri dishes. For fish raised in a naturalistic sensory environment, petri dishes were placed on top of gravel of average size 15 mm [44]. For fish raised in reduced sensory stimulation environment, the petri dishes were placed on plain stainless wire shelves. All fish were placed into their designated sensory environment within 24 h after fertilisation. As a robust way of handling clutch-to-clutch variability for the results shown in Figs 1-6 only one fish from each clutch at each age was assayed. Thus, clutch-to-clutch variability contributed random noise to the data, but no systematic effect.

For the social assay experiments (Fig 7), fish embryos (either WT or *fmr1-/-*) were raised in The University of Queensland aquatic facility until the day before the experiment. Larvae were obtained from 1 L tanks where several males and females were placed together, fed with live rotifers, and used at random without attempting to identify which clutch they came from. The day before imaging about 30 larvae were transported to the lab and kept in a 28.5 °incubator until the imaging session. All test fish were paired with size- and age-matched WT fish. This process was repeated 5 times for each condition and the data combined.

### Hunting behaviour assay

Individual fish were placed into a feeding chamber (CoverWell Imaging Chambers, catalogue number 635031, Grace Biolabs) filled with E3 medium and 30-35 paramecia (*Paramecium caudatum*). The chamber was placed onto a custom made imaging stage consisting of a clear-bottom heating plate at 29.5°C, an infrared LED ring (850 nm, 365 LDR2-100IR2-850-LA powered by PD3-3024-3-PI, Creating Customer Satisfaction (CCS) Inc., Kyoto, Japan) below, and a white LED ring (LDR2-100SW2-LA, CCS) above. Images were recorded using a CMOS camera (Mikrotron 4CXP, Mikrotron) at 500 fps using StreamPix (NorPix, Quebec). Recording of hunting behaviour started after the first attempt for feeding was made by the fish, and each fish was then recorded for 10-15 mins.

### 2-photon calcium imaging

Larvae were embedded in 2.5% low-melting point agarose in the centre of a 35 mm diameter petri dish. Calcium signals in the contralateral tectum to the visual stimulation were recorded with the fish upright using a Zeiss LSM 710 2-photon microscope at the Queenslande Brain Institute’s Advanced Microscopy Facility. Excitation was via a Mai Tai DeepSee Ti:Sapphire laser 463 (Spectra-Physics) at an excitation wavelength of 930-940 nm. Emitted signals were bandpassed (500-550 nm) and detected with a nondescanned detector. Images (416 x 300 pixels) were acquired at 2.2 Hz.

Fish were first imaged for 30 mins in the dark for spontaneous activity (SA). We then recorded tectal responses to stationary 6° diameter dark spots at an elevation of approximately 30° to the fish at either 9 or 11 different horizontal locations (45° to 165° in 15° steps in the first case and 15°to 165° 45 in 15°steps in the second case, where the heading direction of the fish is define as 0°). Only responses to the 9 locations common to all fish were analysed here. Each spot was presented for 1 s followed by 19 s of blank screen a total of 20 times. The presentation order of spot location was randomised, but ensuring that spatially adjacent stimuli were never presented sequentially.

### Social behaviour assay

Custom U-shaped chambers were constructed using a 3D printer. Chambers consisted of 3 compartments separated by 2 glass walls; 2 ‘cue’ compartments each sized 20 × 18 mm and a ‘test’ compartment of length 45 mm (Fig. 7a). Chambers were illuminated using a white LED light strip. A test fish (either WT or *fmr1-/-*) was placed into the test compartment for 5 min to adjust. A WT cue fish was then placed into the left cue compartments. Behavior of both fish was then imaged using a CMOS camera (GrasshopperGS3-U3-23S6M-C, Point Grey) with a 25 mm lens (C-Mount Lens FL-CC2514A-2M, Ricoh) at 100 or 175 fps for 30 mins. For practical reasons (the large number of fish involved and the relatively long rearing time) these fish were raised in featureless dishes.

### Statistical analysis

The Jarque Bera test was used to determine whether data was normally distributed. If any group of data was not normally distributed the Wilcoxon rank sum test was used at each age group to compare effects between genotype. If all groups were normally distributed, ANOVA was used followed by post-hoc t-tests.

## Supplementary Information

### Additional methods

#### Alcian blue staining

Zebrafish larvae were anaesthetised with ethy-3-aminobenzoate (Sigma Aldrich), fixed overnight in 4% PFA/PBS and then washed three times for 10 minutes in PBS. After bleaching in 3% H_2_O_2_/0.5% KOH for 1 hour, larvae were rinsed in 70% ethanol and then stained for 45 minutes using fresh, filtered, alcian blue stain (0.1% alcian blue, 1% HCl, 70% ethanol and 120 mM MgCl_2_). Larvae were washed through 70, 50 and 25% ethanol (all containing 10 mM MgCl_2_) followed by overnight rinse in 25 and 50% glycerol (all with 0.1% KOH). Larvae were mounted in 100% glycerol and photographed with a Zeiss StereoDiscovery V8 microscope and HRc camera using Zen software.

We selected 6 landmarks on the ventral view of the fish and 3 landmarks on the lateral view. In the ventral view, point 1 was defined by the anterior point of Meckel’s cartilage, points 2 and 3 as the posterior most points of the left and right component of Meckel’s cartilage, point 4 as the junction of the left and right components of the ceratohyal cartilage, and points 5 and 6 as the posterior most points of the left and right components of the ceratohyal cartilage. To compare the overall morphological differences between the two genotype, we calculated the pairwise distances bewteen the ventral view landmarks and applied canonical variate analysis (CVA) using MATLAB’s built-in function *canoncorr*. For this computation the genotype variable was represented as binary number, either 0 or 1. The age was rescaled to the range [0,1] so that the canonical coefficients for age and genotype had matching scales and could therefore be directly compared. In lateral views, point 7 was the anterior end of Meckel’s cartilage, point 8 the junction of Meckel’s cartilage and the palatoquadrate, and points 8 and 9 define the lateral axis of the palatoquadrate. Meckel’s cartilage angle (MCA) was measured as the angle between 7-8 and 8-9.

#### Analysis of feeding events

The times at which hunting events began in the recordings were identified manually based on eye convergence [24]. Events were then manually classified based on whether the fish aborted pursuit of the target paramecium (abort event, score 0), pursued but failed to capture the target (miss event, score 1), or the fish successfully captured the target (hit event, score 2 for capture but then eject, 3 for fully capture). Event end was determined by eye deconvergence for abort events, and for other events by the end of the strike bout. The target paramecium was defined as the nearest paramecium towards which the first tuning bout was made.

Automated tracking of the fish and paramecia was performed using custom image processing software in MATLAB as detailed in [27] with minor modifications. In brief, frames were first pre-processed to remove the static background using a Gaussian background model. The approximate location of the fish was identified by connected components analysis on the resulting foreground mask. The position and orientation of the fish were calculated by tracking the midpoint between the eyes and the centre of the swim bladder. This was achieved using a set of correlation filters [74] on pixel values and histogram of oriented gradients features [75]. Filters were rotated through 0,5,10,…,360 degrees and scaled through 60,65,70,…,100% with respect to maximum fish length to accomodate for changes in heading angle and pitch respectively. Filters were trained by manual annotation of the two tracking points in ten randomly selected frames for each fish.

Detection of paramecia was performed using connected components analysis to extract the location of prey-like blobs in each frame from the foreground mask. Multi-object tracking of paramecia between frames was achieved using Kalman filtering and track assignment, which enabled tracking through collisions and short periods of occlusion.

Bout timings and tail kinematics were calculated by first performing morphological thinning and third-order Savitsky-Golay smoothing to extract 101 evenly spaced points along the midline of the tail. Individual bouts were segmented by applying a manually-selected threshold to the amplitude envelope of the mean angular velocity of the most caudal 20% of tail points. Prior to applying the threshold, the angular velocity time series was smoothed using a low-pass filter. The amplitude envelope was estimated using a Hilbert transformation.

From the manual annotations and tracking results, we extracted measures to characterise the hunting efficiency. Abort ratio was calculated as the percentage of aborted events. Hit ratio was calculated as the percentage of events for which the fish successfully captured the prey in its mouth. Inter-bout time was calculated as the averge time between the initiation of feeding related bouts. Detection angle was determined as the angle between the vector defined by the eye midpoint to the target paramecium and the heading angle of the fish.

#### Analysis of neural responses

##### Pre-processing of calcium imaging data

Cell detection and calcium trace extraction were performed using custom MATLAB software as described in [42]. In brief, x-y drifts were corrected using a rigid imaging registration algorithm. Active pixels were identified as pixels that showed changes in brightness over the recording to create an activity map. This activity map was then segmented using a watershed algorithm. For each segmented region, the correlation coeffiecient between pairs of pixels were calcuated. Then, a gaussian mixture model was applied to identify the threshold correlation level for assigning highly correlated pixels to a cell, requiring each cell to contain at least 26 pixels. Once the cells had been identified, we calculated the average brightness of the pixels as the raw fluoresence level *F(t)*. The baseline fluorescence was calculated as a smoothed curve fitted to the lower 20% of the values and the instantaneous baseline level *F*_*0*_*(t)* was taken as the minimum value of the smoothed traced within 3 s centered at t. Neuronal activity levels were calculated as the change of fluoresence level from the baseline as Δ *F/F(t) = (F(t)-F*_*0*_*(t))/F*_*0*_*(t)*. We defined the mean Δ *F/F(t)* over 4 - 7 frames post stimulus presentation as the stimulus-evoked response.

##### Tuning curve

For each neuron, the average responses to each stimulus were averaged to represent the mean response to the given stimulus. We then applied cubic spline interpolation to estimate response amplitute in 5° steps between presented stimuli angles. A Gaussian function was fitted to this interpolated curve to estimate the tuning curve. Neurons with fitted adjusted *R*^*2*^ larger than 0.7 and a maximum evoked response amplitude larger than 1Δ *F/F(t)* were deemed selective neurons and included in further analysis. From the fitted tuning curve, we also obtained the prefered tuning angle and tuning width for each tuned neuron.

##### Assembly properties

Assemblies were detected as detailed previously [42, 76]. In brief, we used a graph theory-based approach to automatically detect assemblies without prior assumptions of expected number of assemblies. For statistical analysis of assembly properties we treated each assembly as a unit. For the area spanned by a given assembly, we first projected all assembly neurons on to the major axis of a fitted ellipse which occupied the NP of the tectum. The normalised distance between the most antieror and postieror assembly members was used to measure the span of the assembly. For assembly tuning, we calculated the mean tuning properties of all neurons belonging to a given assembly.

##### Decoding analysis

To assess how well we could decode the stimulus angle from the responses, we used a Maximum Likelihood decoder (ML) as described in [77]. We assumed that each neuron’s response to a given stimulus *s*_*j*_ was independent, therefore, 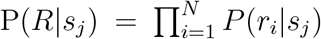. We then estimated the conditional probability that each cell *i* had the response *r*_*i*_ to a given stimulus *s*_*j*_ as P(*r*_*i*_|*s*_*j*_). P(*r*_*i*_|*s*_*j*_) was estimated using the MATLAB *ksdensity* function. The decoded stimulus was the stimulus that gave the highest probability of evoking a given population response, 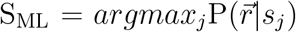. A leave-one-out strategy was used for cross validation: the probablity distributions were estimated with all-but-one trials and we found the stimulus that gives the highest probability to the response that was not included in the estimation, and this process was repeated for each individual trial left out. The decoder performance was calculated as the proportion of correctly identified stimuli out of the 9 stimuli presented. For each stimulus we calculated mean performance, and for each fish we calculated mean performance across all stimuli.

##### Coactivity pattern

To obtain significant coactivity levels we established a threshold using the coactivity patterns during SA. We took the binarised activity pattern and randomly circularly shifted the pattern 1000 times along the time axis, thus preserving the total activity level. The threshold was chosen as the 95th percentile of the shuffled coactivity level. Frames of significant coactivity were collected and divided into different response epochs for further analysis. We applied PCA analysis on the coactivity patterns from different response epochs to quantify the dimensionality of these responses epochs. The similarity between these coactivity pattern was measured by cosine distance. Geometrical relations between EA and SA, SE patterns were measured as the residuals of projections of SA, SE patterns onto the orthonormal basis of EA patterns.

#### Visual environment preference assay

Fish embryos from the same clutch (either WT or *fmr1-/-*) were split into two equally sized groups and reared separately to control for inter-clutch variability across rearing conditions. One group were reared in the naturalistic sensory environment (N) and the other in the reduced sensory stimulation environment (R). Fish were reared until 8 or 9 dpf. Four fish from one of the groups were then placed in a custom circular arena (see below). Free swimming behaviour of the fish was recorded for 20 minutes continuously. Identical imaging was then performed for the second group (*fmr1-/-*: n=20, 20 fish; WT: n=16, 16 fish; for (N) and (R) rearing condition respectively).

The arena was of the same dimensions as the petri dish in which the fish were reared (diameter 85 mm and water depth 5 mm). The arena was made by filling a larger petri dish with 1.2% agrose (UltraPure, Invitrogen) and then cutting a well in the agarose using a 85mm petri dish. A color photographic image of the gravel used for the naturalistic rearing environment, scaled 1:1, was fixed to the underside of one half of the arena. For the other half of the arena we fixed a flat color background which matched the mean hue and brightness of the gravel image (Fig S3a). This image was constructed by randomly shuffling the coordinates of the pixels in the gravel image then smoothing using a 2-dimensional Gaussian filter. The arena was placed onto a custom-made imaging stage illuminated from the side using a stripe of white LED. Images were recorded using a CMOS camera (GrasshopperGS3-U3-23S6M-C, Point Grey) with a 25 mm lens (C-Mount Lens FL-CC2514A-2M, Ricoh), at a rate of 100 fps.

Video data was compressed for convenience using an h264 codec with baseline encoding and quality parameter 17, resulting in visually lossless compression. The position of each fish was tracked using custom software written in MATLAB. The background image was first subtracted by adaptive per-pixel Gaussian modelling on a sliding window comprising every 400th frame spanning a total of 40000 frames (6 minutes and 40 seconds), with a foreground threshold of 2 standard deviations above the mean pixel value. Additionally, a pixel was only considered foreground if its value was above the threshold in at least two of three temporally adjacent frames (the current frame and the two previous frames). Erroneous foreground objects with total area less than 8 pixels were removed using a connected components filter. Remaining foreground object masks were spatially smoothed using a 2-dimensional Gaussian filter and filtered again by connected components to keep only the 4 largest objects which correspond to the four fish. The detected centroids were linked between frames based on minimum Euclidean distance to obtain the trajectory for each fish. We then calculated a gravel preference measure for each fish, defined as the proportion of time that the fish spent on the half of the dish with the gravel substrate.

#### Analysis of social behaviour assay

The locations of the cue and test fish were tracked using custom MATLAB software. Regions of interest (ROIs) were manually drawn for the cue and test chambers respectively to track each fish separately. To model the background a mean image was created using every 500th frame of the movie. To extract a binary image of the fish in each frame, the background was subtracted and pixels with resulting values greater than zero were considered foreground. The location of each fish was computed as the centre of mass of the largest connected component in its corresponding ROI. We calculated the social preference index (SPI) as:

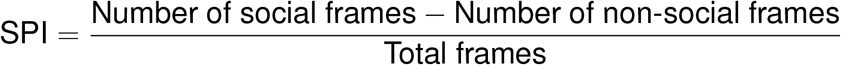

where social frames and non-social frames were defined as a frames for which the test fish was located within the social zone or non-social zone respectively, as shown in Fig. 7a. To quantify the dynamics of fish interaction during social frames we adapted the software written in Python from [47].

For each fish, we calculated the instantaneous speed (mm/s). We considered the cue fish as the reference fish, and identified bout times as the peaks in speed over the full duration of the recording. Peaks were defined as local maxima that were at least two standard deviations greater than the fish’s mean speed. We computed the bout triggered average (BTA) speed of the test fish as the mean over all bouts of the speed of the test fish for the period spanning 200 ms either side of each peak. We quantified the average lag of any movement induced in the test fish by the cue fish as the mean of the delay between each reference peak and the next subseqeunt peak for the test fish. This process was then repeated with the test fish as the reference.

**Figure S1.**
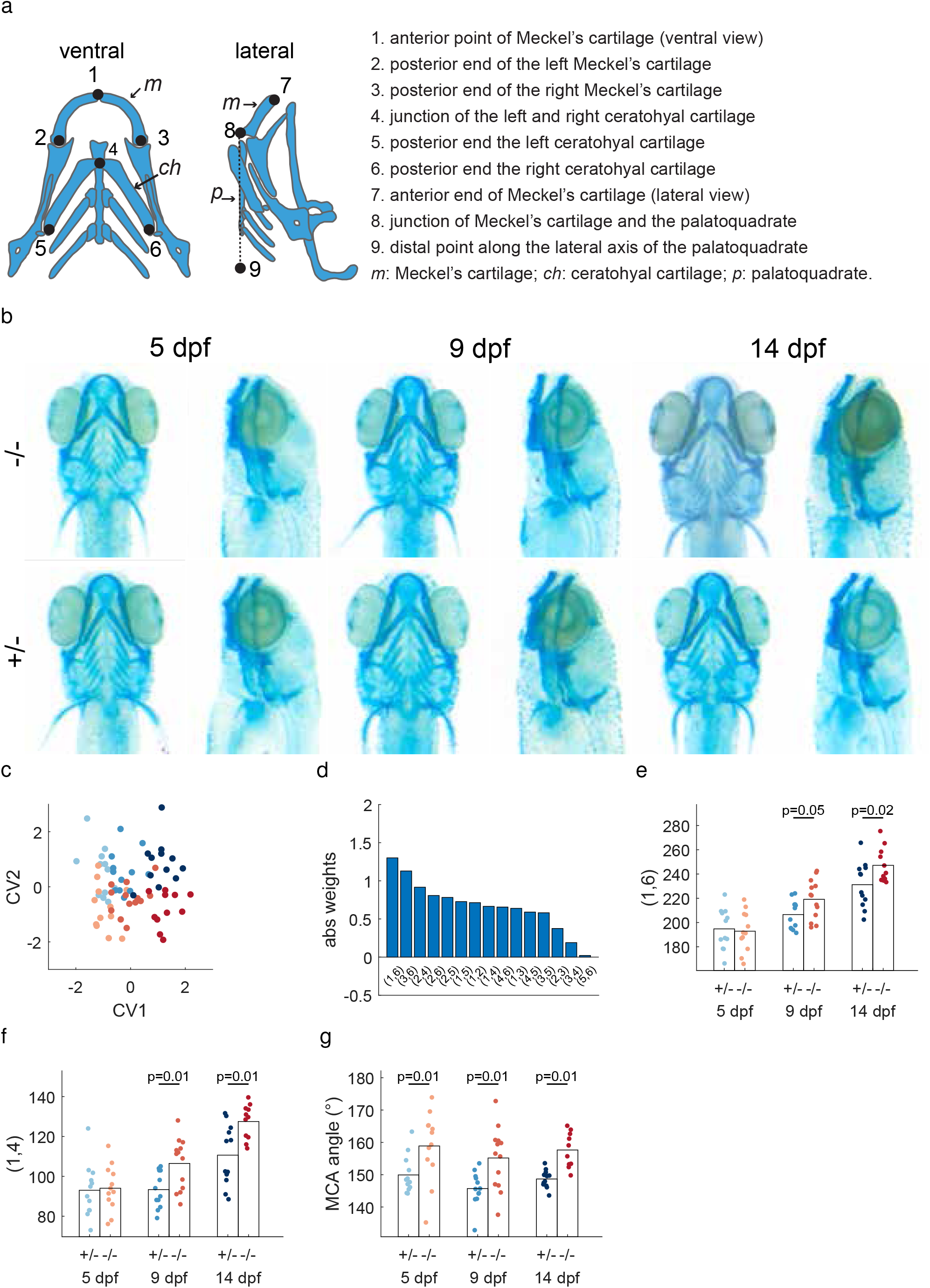
*fmr1-/-* fish show craniofacial abnormalities. **a**. Schematic of the alcian blue-stained cartilages and the landmarks selected for analysis. **b**. Example image of Alcian blue staining of fish at 5,9 and 14 dpf (*fmr1-/-*: n = 12, 12, 12; *fmr1+/-*: n = 12, 13, 10, for each age respectively). **c**. CVA analysis revealed significant association between morphological traits and the age and genotype of the fish. CV1 reflects correlation with age (p = 10^−15^; magnitude of canonical coefficients *b*_CV1,age_ = 2.45 and *b*_CV1,genotype_ = 0.20; See Extended methods). CV2 reflects correlation with genotype (p = 10^−4^; *b*_CV2,age_ = 0.20 and *b*_CV2,genotype_ = 1.98). **d**. The magnitude of the weights of CV2 for different distances between the landmarks on the ventral view. **e**. The distance with the highest weight, (1, 6), was larger in *fmr1-/-* fish at 9 and 14 dpf. **f**. The distance (1, 4), equivalent to lower jaw length, was larger in *fmr1-/-* fish at 9 and 14 dpf. **g**. The Meckel’s cartilage angle (MCA, between points 7,8, and 9) was less acute in *fmr1-/-* fish.

**Figure S2.**
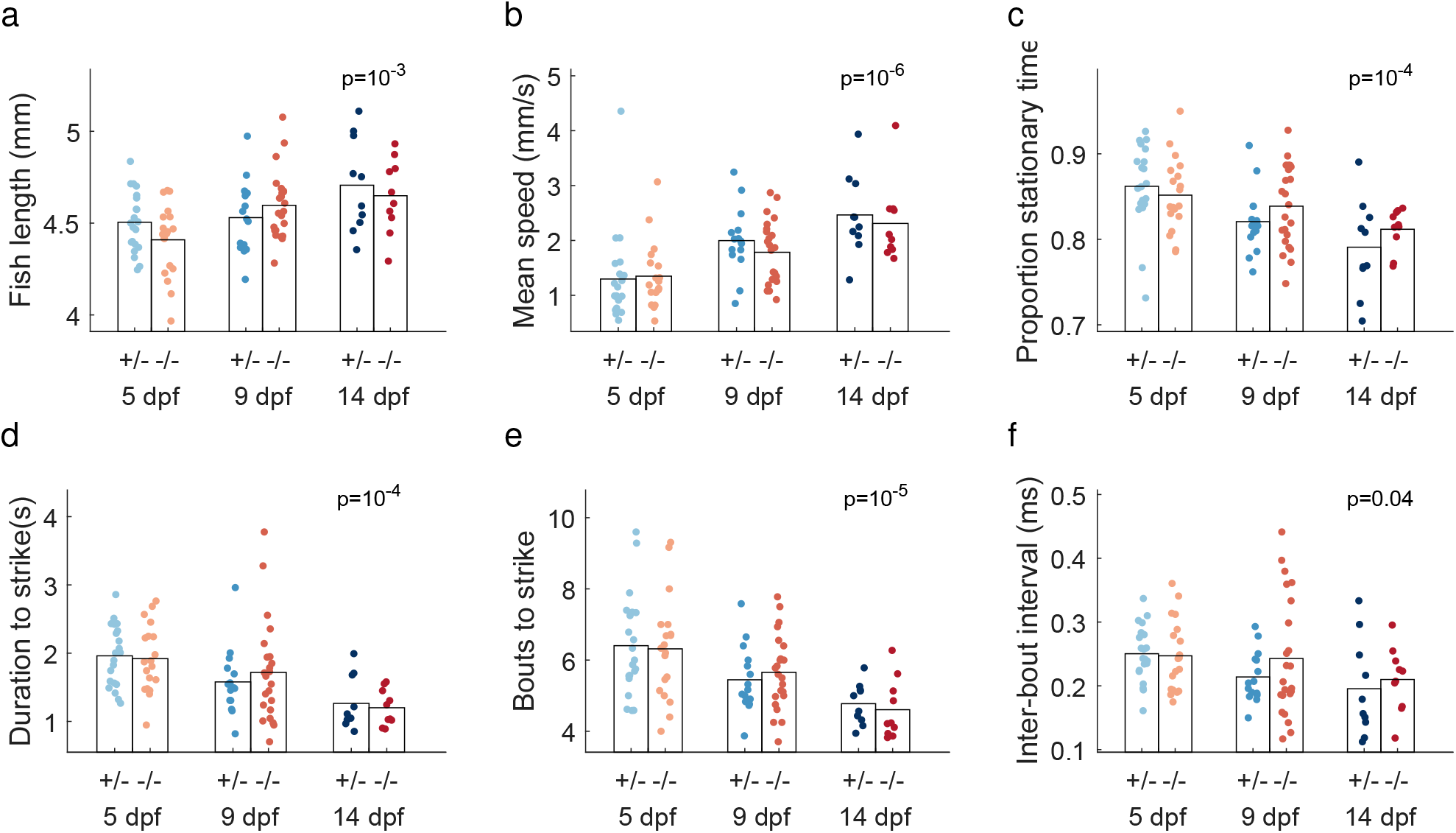
*fmr1-/-* fish did not show any motor deficits during hunting. **a-c**. Fish length, mean swimming speed, and the proportion of stationary time during hunting was similar between genotypes. **d-f**. The duration to strike, the number of bouts made before a strike and the inter-bout interval during a hunting sequence were not different between genotypes. All measures showed significant differences with age (p values show age effect from one-way ANOVA), indicating a development trend of more efficient hunting over age.

**Figure S3.**
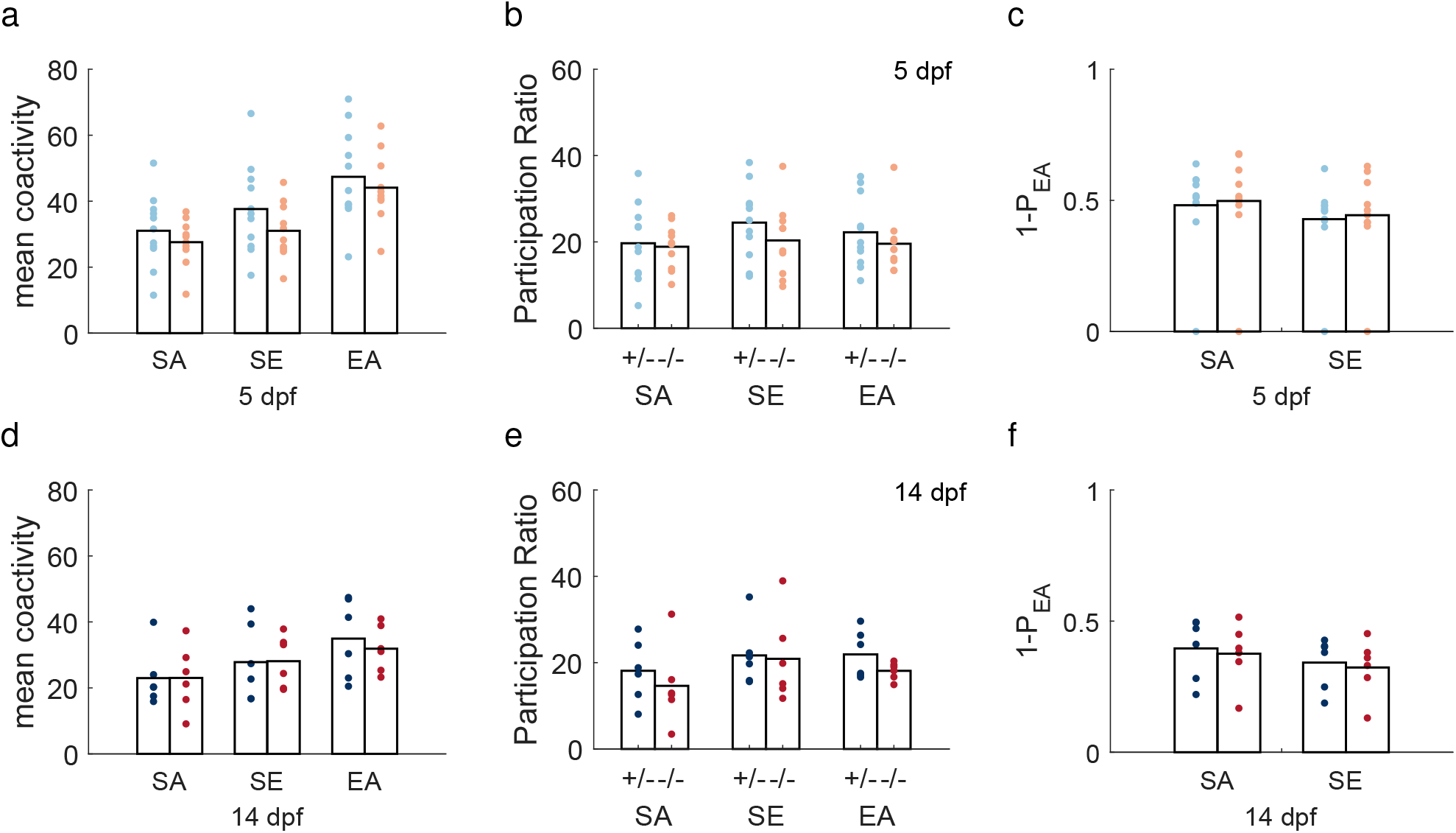
Tectal coactivity patterns were not altered in *fmr1-/-* fish at 5 and 14 dpf. **a-c**. Mean coactivity level, participation ratio and residuals of SA and SE patterns on EA patterns at 5 dpf. **d-f**. Same measures at 14 dpf. There were no significant differences between genotypes.

**Figure S4.**
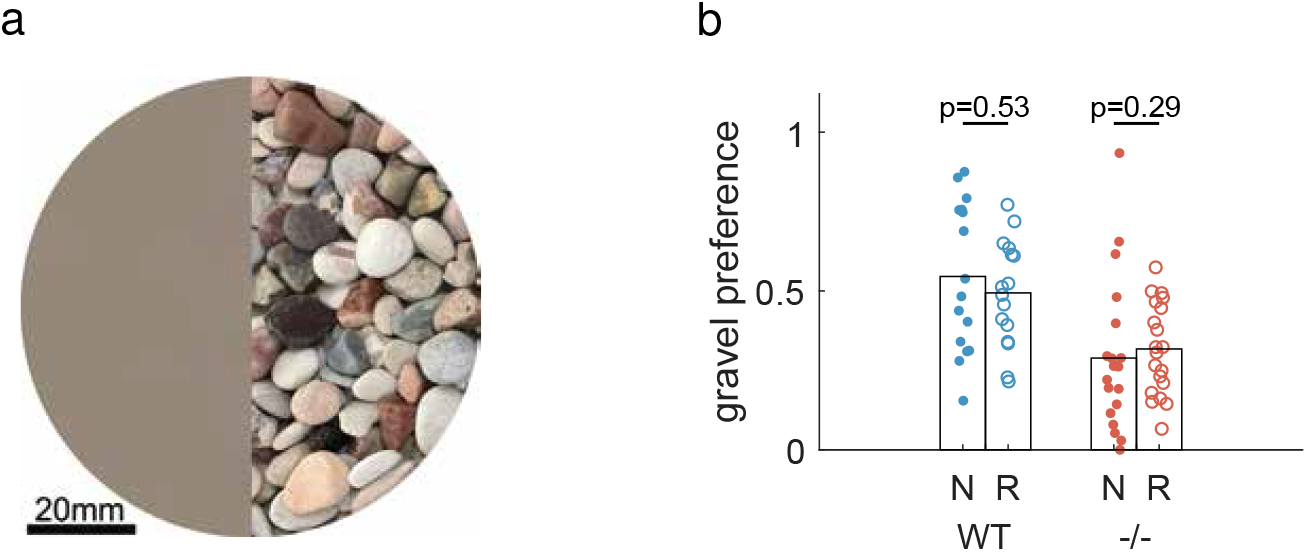
Gravel preference was independent of rearing condition. **a**. The image placed underneath the dish in which the fish were swimming. The featureless side (left) of the image was produced by scrambling and smoothing the gravel image (right) to ensure average brightness and color are matched (See Additional methods). **b**. Rearing condition did not affect the gravel preference of either genotype.

